# Differential effects of variations in human P450 oxidoreductase on the aromatase activity of CYP19A1 polymorphisms R264C and R264H

**DOI:** 10.1101/664839

**Authors:** Shaheena Parween, Giovanna DiNardo, Francesca Baj, Chao Zhang, Gianfranco Gilardi, Amit V. Pandey

**Affiliations:** Pediatric Endocrinology, Diabetology, and Metabolism, Department of Pediatrics, University Children’s Hospital Bern, 3010, Bern, Switzerland; Department of Biomedical Research, University of Bern, 3010 Bern, Switzerland; Department of Life Sciences and Systems Biology, University of Torino, Via Accademia Albertina 13, Torino, Italy

**Keywords:** CYP19A1, Aromatase, P450 oxidoreductase, estrogen synthase, POR, cytochrome P450

## Abstract

Aromatase (CYP19A1) converts androgens into estrogens and is required for female sexual development and growth and development in both sexes. CYP19A1 is a member of cytochrome P450 family of heme-thiolate monooxygenases located in the endoplasmic reticulum and depends on reducing equivalents from the reduced nicotinamide adenine dinucleotide phosphate via the cytochrome P450 oxidoreductase coded by *POR*. Both the *CYP19A1* and *POR* genes are highly polymorphic, and mutations in both these genes are linked to disorders of steroid biosynthesis. We have previously shown that R264C and R264H mutations in *CYP19A1*, as well as mutations in *POR*, result in a reduction of CYP19A1 activity. The R264C is a common polymorphic variant of *CYP19A1*, with high frequency in Asian and African populations. Polymorphic alleles of *POR* are found in all populations studied so far and, therefore, may influence activities of *CYP19A1* allelic variants. So far, effects of variations in *POR* on enzymatic activities of allelic variants of *CYP19A1* or any other steroid metabolizing cytochrome P450 proteins have not been studied. Here we are reporting the effects of three POR variants on the aromatase activities of two CYP19A1 variants, R264C and R264H. We used bacterially expressed and purified preparations of WT and variant forms of CYP19A1 and POR and constructed liposomes with embedded CYP19A1 and POR proteins and assayed the CYP19A1 activities using radiolabeled androstenedione as a substrate. With the WT-POR as a redox partner, the R264C-CYP19A1 showed only 15% of aromatase activity, but the R264H had 87% of aromatase activity compared to WT-CYP19A1. With P284L-POR as a redox partner, R264C-CYP19A1 lost all activity but retained 6.7% of activity when P284T-POR was used as a redox partner. The R264H-CYP19A1 showed low activities with both the POR-P284L as well as the POR-P284T. When the POR-Y607C was used as a redox partner, the R264C-CYP19A1 retained around 5% of CYP19A1 activity. Remarkably, The R264H-CYP19A1 had more than three-fold higher activity compared to WT-CYP19A1 when the POR-Y607C was used as the redox partner, pointing towards a beneficial effect. The slight increase in activity of R264C-CYP19A1 with the P284T-POR and the three-fold increase in activity of the R264H-CYP19A1 with the Y607C-POR point towards a conformational effect and role of protein-protein interaction governed by the R264C and R264H substitutions in the CYP19A1 as well as P284L, P284T and Y607C variants of POR. These studies demonstrate that the allelic variants of P450 when present with a variant form of POR may show different activities, and combined effects of variations in both the P450 enzymes as well as in the POR should be considered when genetic data are available. Recent trends in the whole-exome and whole-genome sequencing as diagnostic tools will permit combined evaluation of variations in multiple genes that are interdependent and may guide treatment options by adjusting therapeutic interventions based on laboratory analysis.

## 1 Introduction

Aromatase (CYP19A1, EC: 1.14.14.14) regulates estrogen biosynthesis [1] by converting androgens into estrogens [2, 3]. CYP19A1 is a 503 amino acid protein (NP_000094) encoded by the *CYP19A1* gene (GeneID:1588, NCBI: NM_000103), located on chromosome 15 (15q21.2, GRCh38 15:51208056-51338597). CYP19A1 is a membrane-bound protein located in the endoplasmic reticulum (ER) and is a member of the cytochrome P450 superfamily. Cytochrome P450 enzymes are involved in the biosynthesis of steroid hormones, and they also metabolize drugs and xenobiotics [4]. Two different types of the cytochrome P450 family of proteins are present in humans [5]. The P450 type 1 proteins are found in the mitochondrion and are responsible for the metabolism of steroid hormones and sterols and are targets for drugs in a range of disorders [6-11]. The P450 type 2 proteins are localized in the endoplasmic reticulum and have a range of metabolic activities, including drugs, xenobiotics as well as endogenous substrates, and steroid hormones like progesterone, dehydroepiandrosterone, and testosterone [12-17]. Genetic defects in steroid metabolizing cytochromes P450 and their redox partners cause metabolic disorders with disordered steroidogenesis [18-25]. CYP19A1 is highly expressed in ovaries and plays a significant role in the regulation of the reproductive cycle in females [26]. In males, CYP19A1 is expressed in the gonads, and the paracrine effects of CYP19A1 reaction products are required for normal spermatogenesis [27-30].

Additionally, CYP19A1 expression is also found in extra-gonadal tissues, including liver, muscle, placenta, bone, breast, adipose tissue, and brain [26, 31, 32]. Initial work on the characterization of CYP19A1 was done on enzyme preparations purified from the placenta by laboratories of Hall [33], Vickery [34], and Harada [35] and a placental protein preparation was also used for the determination of X-ray crystal structure of CYP19A1 protein by Ghosh [36]. The CYP19A1 protein is required for the biosynthesis of estrogens (C18 steroids) from the androgen precursors (C19 steroids) [1, 37]. Some of the critical reactions catalyzed by CYP19A1 include the biosynthesis of estrone (E1) from androstenedione, estriol (E3) from 16-hydroxytestosterone and 17β-estradiol (E2) from testosterone (Figure 1) [1]. The androstenedione and testosterone are well known as the most common physiological steroid substrates for CYP19A1 [1, 38, 39]. Recently 16β-OH-androstenedione was also reported to be a substrate for CYP19A1 [40]. The catalytic process of aromatization of androgens is multifaceted, and it comprises of three enzymatic reaction steps involving the transfer of three pairs of electrons from the co-factor reduced nicotinamide adenine dinucleotide phosphate (NADPH) and consumption of oxygen. Electron transfer from NADPH to CYP19A1 is carried out by NADPH cytochrome P450 oxidoreductase (POR) [41, 42]. The initial two steps of aromatase reaction are two sequential hydroxylation reactions at the 19-methyl group of androgens, whereas the last step causes the aromatization of the A-ring of the steroid, which is specific for CYP19A1 [43-45].

**Figure 1:**
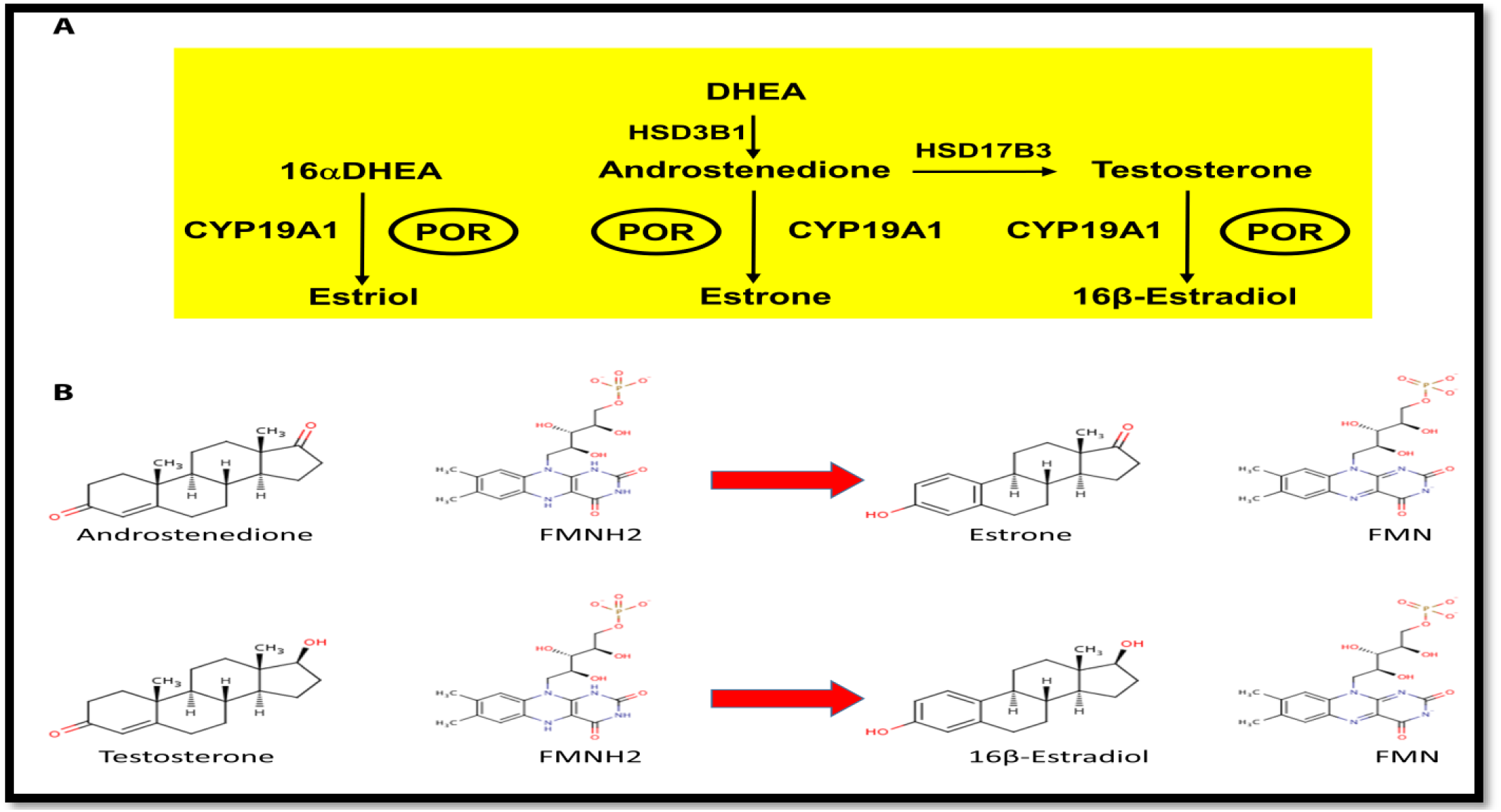
Steroid hormone biosynthesis and role of POR. **A:** In the human placenta, DHEA (CHEBI:28689) is metabolized to form androstenedione. Androstenedione then serves as a substrate for the formation of testosterone (CHEBI:17347) which is then converted into estrogens. In the liver of the fetus, DHEAS (CHEBI:16814) is metabolized into 16αDHEAS (CHEBI:87774), which then gets converted into 16αDHEA inside the placenta to form estriol (CHEBI:27978). The biosynthesis of estrogens requires the aromatase activity of CYP19A1, which is reliant on POR. Figure adapted from our previous publication by Udhane et al. 2017 [64]. **B:** Electron transfer from NADPH to CYP19A1 is carried out via POR through reduced FAD and FMN. Chemical structures were obtained from ChEBI (https://www.ebi.ac.uk/chebi/).

Multiple genetic defects are associated with the reduced CYP19A1 activity in humans (OMIM: 107910, 613546, 139300) [19, 46]. In females, a reduction in CYP19A1 activity causes abnormal external genitalia in the female fetus, maternal virilization during gestation, healthy development of ovaries but the increased occurrence of polycystic ovary syndrome, and virilization at puberty [31, 38]. The first case of CYP19A1 deficiency in humans was described by Shozu *et al*. [2]. Later Morishima *et al*. described that CYP19A1 deficiency occurs during fetal life in both males and females [47]. Effect of CYP19A1 deficiency on the maturation of skeleton, growth, and bone homeostasis [27, 48], as well as abnormal plasma lipid profile and variations in insulin resistance, have been reported in multiple publications [19, 30, 31, 38, 49-52]. Estrogen excess has also been shown in many publications to cause an increase in the expression and activity of CYP19A1 [53, 54]. Several natural compounds, including nicotine, vitamin E, resveratrol and curcumin, may inhibit CYP19A1 activity [8, 55, 56]. CYP19A1 activity is also inhibited by bisphenol A and sildenafil [55]. CYP19A1 inhibitors like anastrozole are used in the treatment of breast cancer and to treat the excess production of estrogen in men [57].

POR deficiency (PORD, OMIM: 613537 and 201750), is a rare form of congenital adrenal hyperplasia, that was first reported in patients with disordered steroidogenesis [24], and later on, multiple disorders have been linked to mutations in POR [42, 58-61]. While many cytochromes P450 have been tested with different POR variants, allelic variants of the steroid metabolizing cytochromes P450 have not been studied for the potential changes in enzyme activities caused by variant forms of POR [42, 60, 62]. Subramanian *et al*. had examined the combined effect of P450 and POR variations in a report describing activities of three common alleles of CYP2C9 assayed with the WT and variant forms of POR [63]. Since POR is obligate electron donor for all cytochrome P450 proteins in the endoplasmic reticulum, variations in POR should also be taken into account when studying the effect of genetic polymorphisms in cytochromes P450. Loss of CYP19A1 activity may severely affect the production of estrogens, and several variations of POR exist with significant population distribution, which can alter enzymatic activities of CYP19A1 [64-68]. We, therefore, tested the impact of some population variants of POR on estrogen formation by the two variants of CYP19A1. The R264C (<u>rs700519</u>) is a common polymorphic variant of CYP19A1(CYP19A1*4) while the R264H (<u>rs2304462</u>) is a rare variant. We generated the WT and variant forms of human POR and CYP19A1 in bacteria and used lipid-based reconstitution to do the enzymatic analysis of CYP19A1-aromatase activity using androstenedione as a substrate. We also studied the effect of CYP19A1 variations on protein stability by limited proteolysis to understand the potential causes of the changes in enzyme activities by different variants. This study shows that variations in POR may have variable effects on activities of CYP19A1 variants, potentially leading to altered steroid metabolism in individuals with variant CYP19A1 alleles.

## 2 Materials and methods

### Materials

Tris-base, NADPH, acetic acid, magnesium acetate, Sucrose, potassium phosphate EDTA, DTT, glycerol, PMSF, and Benzonase were purchased from Sigma-Aldrich Chemie GmbH (Buchs, Switzerland). Carbenicillin, FeCl_3_, ZnCl_2_, CoCl_2_, Na_2_MoO_4_, CaCl_2_, CuCl_2_, M H_3_BO_3_ were purchased from CarlRoth GmBH (Switzerland). Goat anti-rabbit antibodies labeled with infra-red dyes were from LI-COR Bioscience Inc. (NE, USA). The protein assay dye reagent (RC-DC assay) was from Bio-Rad (Hercules, CA). The anti-POR antibody was from Genscript (NJ, USA).

### Recombinant expression and purification of CYP19A1 and its polymorphic variants R264H and R264C

The experiments were carried out with a well-characterized recombinant soluble human CYP19A1, which has the structure and activity comparable to full-length CYP19A1 [69]. The polymorphic variants R264H and R264C of CYP19A1 were generated as described previously by Baravalle *et al*. in 2017 [70]. The WT CYP19A1 and the two polymorphic variants, R264H and R264C, were expressed and purified based on previously described methods [69, 71-73]. Briefly, the *E. coli* DH5α rubidium-competent cells were transformed with a pCWOri+ plasmid vector carrying the *CYP19A1* gene and selected with 100 μg/mL ampicillin. The CYP19A1 protein expression was induced by the addition of 1 mM Isopropyl β-D-1-thiogalactopyranoside. Liquid cultures were grown in the Terrific Broth medium in the presence of the heme precursor δ-aminolevulinic acid at 28°C for 48 hours. Bacterial cells were collected and re-suspended in 100 mM potassium phosphate (pH 7.4) containing 20 % glycerol, 1 % Tween-20, and 1 mM β-mercaptoethanol supplemented with 1 mg/mL lysozyme and a protease inhibitor cocktail (Roche, Switzerland) at 4°C. Bacteria were broken by sonication and ultra-centrifuged at 100000 x g for 20 minutes. The cellular supernatant containing the soluble proteins was loaded on an ion-exchange column (DEAE-Sepharose Fast-Flow, GE Healthcare), followed by processing over a Nickel-ion Chelating-Sepharose Fast-Flow affinity column (GE Healthcare). The CYP19A1 protein was eluted from the Nickel-Sepharose column with a gradient of 1 mM to 40 mM histidine, which was later removed by ultrafiltration on the 30 kD molecular weight cut-off Amicon concentrator device (Merck-Millipore).

### POR expression and membrane purification

The WT POR and variants were produced in bacteria using the plasmid constructs described in earlier publications [68, 74-76]. The protocol for the expression of POR variants and membrane as well as liposome preparation is reported in our earlier publications [64-66, 68, 74, 77].

### CYP19A1 activity measurement in the reconstituted liposome system

CYP19A1 activity of proteins was monitored by the tritiated water release assay based on an earlier method described by Lephart and Simpson [78] with slight modifications that have been described previously using an assay system containing lipids, POR and CYP19A1 [64, 68]. Androstenedione was used as the substrate for the CYP19A1 reaction. POR and CYP19A1 proteins were mixed into liposomes containing 1,2-Dilauroyl-sn-glycero-3-phosphorylcholine (DLPC) and 1,2-Dilauroyl-sn-glycero-3-phosphorylglycerol (DLPG). Reaction mixture consisted of 50 pmol of CYP19A1, 200 pmol of POR and 20,000 cpm of ^3^H androstenedione ([1^β_3^H(N)]-andros-tene-3,17-dione;) in 100 mM potassium phosphate buffer (pH 7.4) with 100 mM NaCl. Different concentrations of androstenedione (0.01–30 µM) were used for assays to study the enzyme kinetics. The enzymatic reaction of CYP19A1 was started by adding NADPH to 1 mM concentration, and reaction mixtures were incubated with shaking for 1 h. The reaction was stopped by adding chloroform and the water phase was mixed with a 5% charcoal/0.5% dextran solution and centrifuged at 14000 x g for 5 minutes and 0.5 ml of supernatant was used for ^3^H radioactivity counting. The specific activity of human CYP19A1 was calculated as (pmol of product/minute of incubation/nmol of P450), where pmol of the product formations were calculated as:

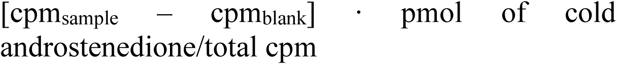

Data were analyzed using GraphPad Prism (GraphPad, La Jolla, CA) based on Michaelis-Menten kinetics to determine the Vmax and Km. Values obtained from triplicate experiments were expressed as mean ± S.E.M. (standard error of mean).

### FASTpp assay of CYP19A1 and its polymorphic variants R264H and R264C

The FASTpp assay was performed as described previously [66, 79, 80]. Reaction mixtures contained 0.2 mg/mL of CYP19A1 WT or polymorphic variants R264H and R264C, 10 mM CaCl_2_, 0.05 mg/mL thermolysin in 0.1 M potassium phosphate buffer (pH 7.4) containing 20% glycerol, 0.1 % Tween-20, and 1 mM β-mercaptoethanol. A negative control reaction was carried out without including thermolysin. Protein digestion was performed for one minute in a TGradient Thermoblock (Biometra, Switzerland), generating a temperature gradient from 30 to 70°C, and reactions were stopped by addition of 10 mM EDTA (pH 8.0). Samples were analyzed by SDS-PAGE and quantified using Image Lab software (Bio-Rad, Hercules, CA).

## 3 Results

### Population Genetics of CYP19A1 amino acid variants R264C and R264H

The R264C variant of CYP19A1 is present at a low frequency in the European population with a minor allele frequency (MAF) of 0.02 in the 1000 Genomes database (Table 1). Among the American population, the MAF of R264C was 0.04. In the East Asian (MAF 0.17), South East Asian (MAF 0.24), and African (MAF 0.18) populations, the R264C variant was present at a much higher frequency. Looking individually, the Italian population from Tuscany had the lowest occurrence of R264C with **a** MAF of 0.006 in the HapMap database (Table 1). In the HapMap database, the R264C variant was found in a Mexican population with a MAF of 0.01, while the Japanese population showed the highest presence of R264C with a MAF of 0.27. HapMap data showed similar MAF among subpopulations as reported in 1000 Genome database, the only exception being the Maasai population from Kenya, which had the R264C allele present at the MAF of 0.04, which is much lower than the distribution seen in other African populations. On the other hand, the R264H variant of CYP19A1 (rs2304462) is found only as a rare variant present at a MAF of 0.00003 in ExAc Aggregated Populations and at a MAF of 0.00002 in gnomAD-Exomes.

**Table 1:** Genetic data for the CYP19A1 variations. The R264C (DBSNP: rs700519) is a common polymorphism in the CYP19A1 gene. Population codes referred to in the table are for **1000 Genome**: **AMR** American; **AFR** African; **EAS** East Asian; **SAS** South East Asian; **EUR** European. For the **HapMap** project: **ASW**, African ancestry in Southwest USA; **CEU**, Utah residents with Northern and Western European ancestry; **CHB**, Han Chinese in Beijing, China; **CHD**, Chinese in Metropolitan Denver, Colorado; **GIH**, Gujarati Indians in Houston, Texas; **JPT**, Japanese in Tokyo, Japan; **LWK**, Luhya in Webuye, Kenya; **MEX**, Mexican ancestry in Los Angeles, California, USA; **MKK**, Maasai in Kinyawa, Kenya; **TSI**, Toscans in Italy; **YRI**, Yoruba in Ibadan, Nigeria. The R264H (rs2304462) is a rare variant and has been seen with lower frequencies across populations. Data for population genetic distribution of variants were retrieved from the NCBI (http://www.ncbi.nlm.nih.gov). Populations with a minor allele frequency that is significantly higher than average are shown in bold.

| Population ID-Class | Number of alleles | Major Allele Freq. | Minor Allele Freq. |
| --- | --- | --- | --- |
| **R264C (rs700519 )** |  |  |  |
| **1000 Genome** | 5008 | G=0.860 | A=0.140 |
| 1000 Genome- AMR | 694 | G=0.96109998 | A=0.03890000 |
| 1000 Genome-AFR | 1322 | G=0.81770003 | A=0.18230000 |
| 1000 Genome-EAS | 1008 | G=0.82940000 | A=0.17060000 |
| **1000 Genome-SAS** | 978 | G=0.76279998 | **A=0.23720001** |
| 1000 Genome-EUR | 1006 | G=0.97119999 | A=0.02880000 |
| **ESP-cohort North America** | 4542 | G=0.91391456 | A=0.08608542 |
| **HapMap** |  |  |  |
| <u>HapMap-CEU</u> | 224 | G=0.96875000 | A=0.03125000 |
| <u>HapMap-TSI</u> | 174 | G=0.99425286 | A=0.00574713 |
| <u>HapMap-MEX</u> | 98 | G=0.98979592 | A=0.01020408 |
| <u>HapMap-CHD</u> | 170 | G=0.88235295 | A=0.11764706 |
| **<u>HapMap-JPT</u>** | 172 | G=0.72674417 | **A=0.27325583** |
| <u>HapMap-ASW</u> | 98 | G=0.83673471 | A=0.16326530 |
| <u>HapMap-GIH</u> | 174 | G=0.83908045 | A=0.16091955 |
| <u>HapMap-CHB</u> | 82 | G=0.85365856 | A=0.14634146 |
| **<u>HapMap-YRI</u>** | 224 | G=0.78571427 | **A=0.21428572** |
| <u>HapMap-LWK</u> | 180 | G=0.88888890 | A=0.11111111 |
| <u>HapMap-MKK</u> | 286 | G=0.95804197 | A=0.04195804 |
| R264H (rs2304462) |  |  |  |
| **ExAc Aggregated Populations** | 121366 | C=0.99996704 | T=0.00003296 |
| **gnomAD-Exomes** | 246062 | C=0.99998 | T=0.00002 |

### Genetics of variants in POR

The POR variants described in this study were selected by searching the genetic sequence databases. Selected variants were subjected to stability analysis on protein structure followed by sequence conservation studies to filter the variants with disease-causing possibilities [59]. Human POR protein has 680 amino acids and has evolved from the ferredoxin reductase and ferredoxin like structures to create a single P450 reductase protein in higher eukaryotes [81]. The POR variations described here are highly conserved among different species and are described as rare genetic variants in the Exome database of NCBI (Table 2).

**Table 2:** Minor allele Frequencies of variants in POR described in this study according to the Exome database of NCBI.

| POR | DBSNP id | Global | European | Asian | American | African |
| --- | --- | --- | --- | --- | --- | --- |
| **P284L** | rs72557938 | T=0.00005 | T=0.00004 | T=0.0001 | T=0.0000 | T=0.0000 |
| **P284T** | rs72557937 | A=0.00004 | A=0.00000 | A=0.0000 | A=0.0000 | A=0.0005 |
| **Y607C** | rs72557954 | G=0.00011 | G=0.00008 | G=0.0003 | G=0.0000 | G=0.0000 |

The P284L is a rare variant found mostly in Europeans, and the P284T variant had been only observed in an African population [64, 82, 83]. The Y607C is a rare genetic variant found in both the Asian and European populations (Table 2).

### Structure analysis of variants in CYP19A1 and POR

The R264 residue on CYP19A1 is surface exposed and located on the G helix of the CYP19A1 structure (Figure 2A), which is part of the substrate access channel, and it has been widely reported to be flexible in cytochromes P450. R264 is the central residue of a 3 Arg-containing cluster that is known as a consensus sequence for multiple kinases. Thus, it can play a central role in both the formation of substrate access and CYP19A1 activity by post-translational modifications (51). The amino acid P284 is situated in the flexible hinge region, between the FAD and FMN binding domains of POR (Figure 2B). This hinge region is responsible for flexibility and domain movements in POR. Mutations in the POR that are located in the hinge region may cause conformational changes that would change the interactions between POR and its protein/small molecule partners [42, 64]. The Y607 residue is located close to the NADPH binding site (Figure 2B), and the Y607C variant in POR has been shown to reduce the CYP19A1 catalytic efficiency by 90% [65].

**Figure 2:**
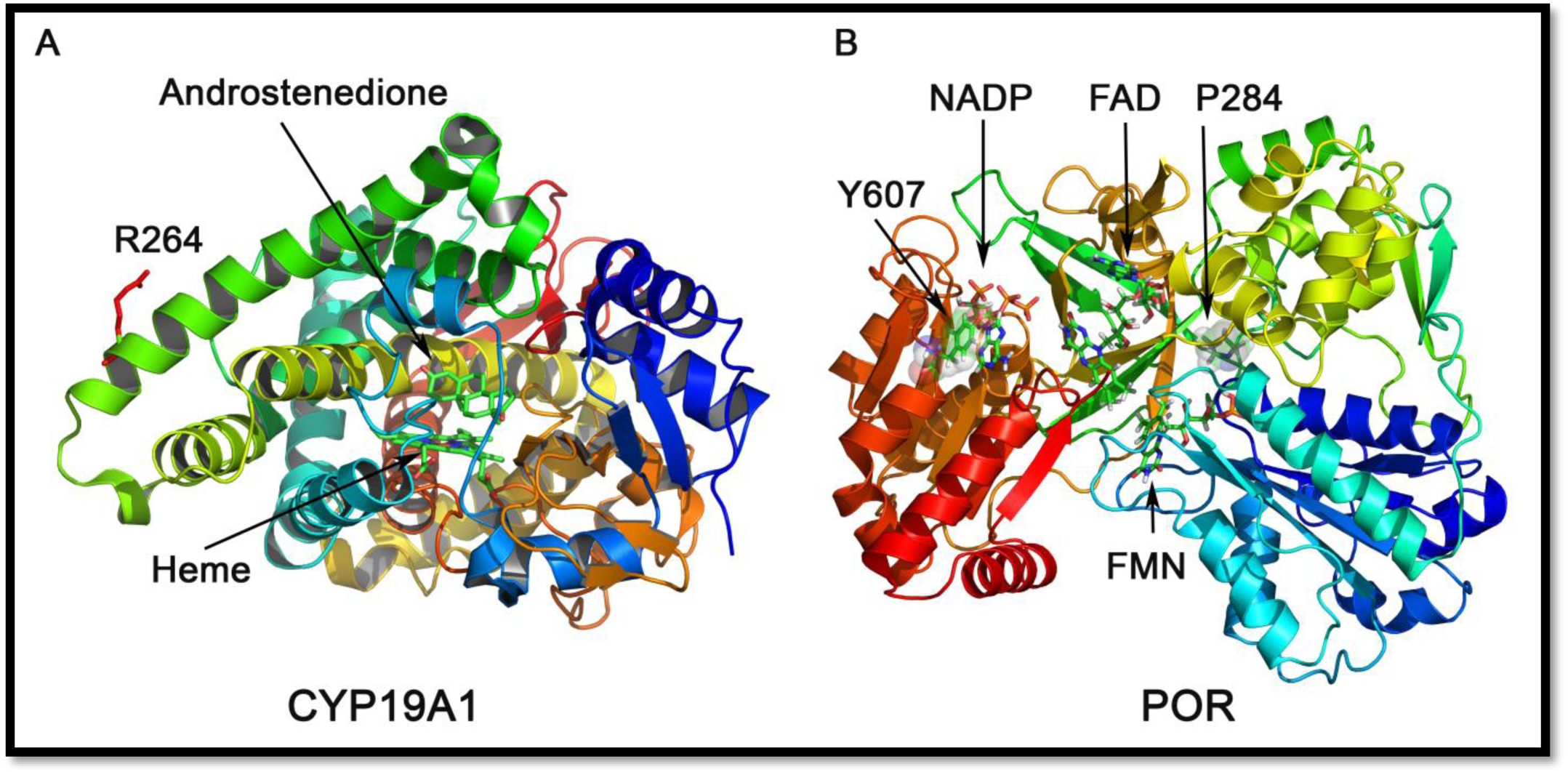
Location of CYP19A1 and POR variants: **A**. Locations of CYP19A1 variants in protein structure. The R264 residue on CYP19A1 is surface exposed and located on the G helix of the CYP19A1 structure, which is part of the substrate access channel, and it has been widely reported to be flexible in cytochromes P450. Here a ribbons model of CYP19A1 protein is shown, and the bound substrate, androstenedione, is depicted as sticks model. **B**. Locations of POR variants. X-ray crystal structure of the human POR (**PDB#3QE2**) depicting the positions of POR variations described in this study. The co-factors FAD, FMN, and NADP are depicted as sticks. The P284 residue is situated in the hinge region of the POR. The hinge region of POR is vital for flexibility and conformation shifts in POR, and mutations in this region of POR may create changes in conformation that would modify the interactions of POR with its partner proteins. The Y607 residue is located in the NADPH binding site of POR. Binding of NADPH is required for the transfer of electrons to redox partners of POR as well as direct reduction of dyes and small molecules by POR.

### Enzyme activity of CYP19A1 and its polymorphic variants R264H and R264C supported by POR-WT

We performed *in-vitro* CYP19A1 activity assays with the WT CYP19A1 and its polymorphic variants (R264H and R264C) to study the effects of the two variations on the enzyme activity. To this aim, reconstituted liposomes were prepared by mixing the purified recombinant CYP19A1 with purified membranes containing WT-POR as described in materials and methods. Then we assessed the enzyme activity by standard tritiated water release assay. For the R264H variant of CYP19A1, the apparent Km for androstenedione was comparable to that of WT CYP19A1, whereas the apparent Vmax was reduced by ∼30% (Figure 3, Table 3). The catalytic efficiency of the CYP19A1-R264H variant was reduced by ∼13% compared to WT CYP19A1. However, for the R264C variant of CYP19A1, the apparent Km for androstenedione increased by 3.2 fold as compared to WT CYP19A1. The apparent Vmax was reduced by ∼50%.

**Table 3.** Kinetic parameters for activities of CYP19A1 and its polymorphic variants supported by WT POR. Variable androstenedione concentration (0.3–30 µM) were used for the biotransformation of androstenedione into estrone. All assays were performed with 1 mM NADPH. Data are shown as mean ± SEM of independent replicates (n=3). Catalytic efficiencies of different combinations of POR and CYP19A1 variants are given as a percentage of WT-POR and WT-CYP19A1 combination (% of WT-WT catalytic efficiency). * indicates the significance based on two tailed students t-test (*p<0.05).

| | Km Androstenedione<br>(μM) | Vmax<br>pmol/min/nmol | Vmax/Km | % of WT-WT<br>catalytic<br>efficiency |
| --- | --- | --- | --- | --- |
| ***CYP19A1 (aromatase) activity for the Conversion of androstenedione to estrone*** |  |  |  |  |
| CYP19A1-WT | 1.6 ± 0.9 | 35.5 ± 4.7 | 21.6 | 100.0 |
| CYP19A1-R264H | 1.3 ± 0.9 | 25.2 ± 4.1 | 18.8 | 87.0 |
| CYP19A1-R264C | 5.1 ± 2.6 | 16.9 ± 2.8* | 3.3* | 15.3* |

**Figure 3:**
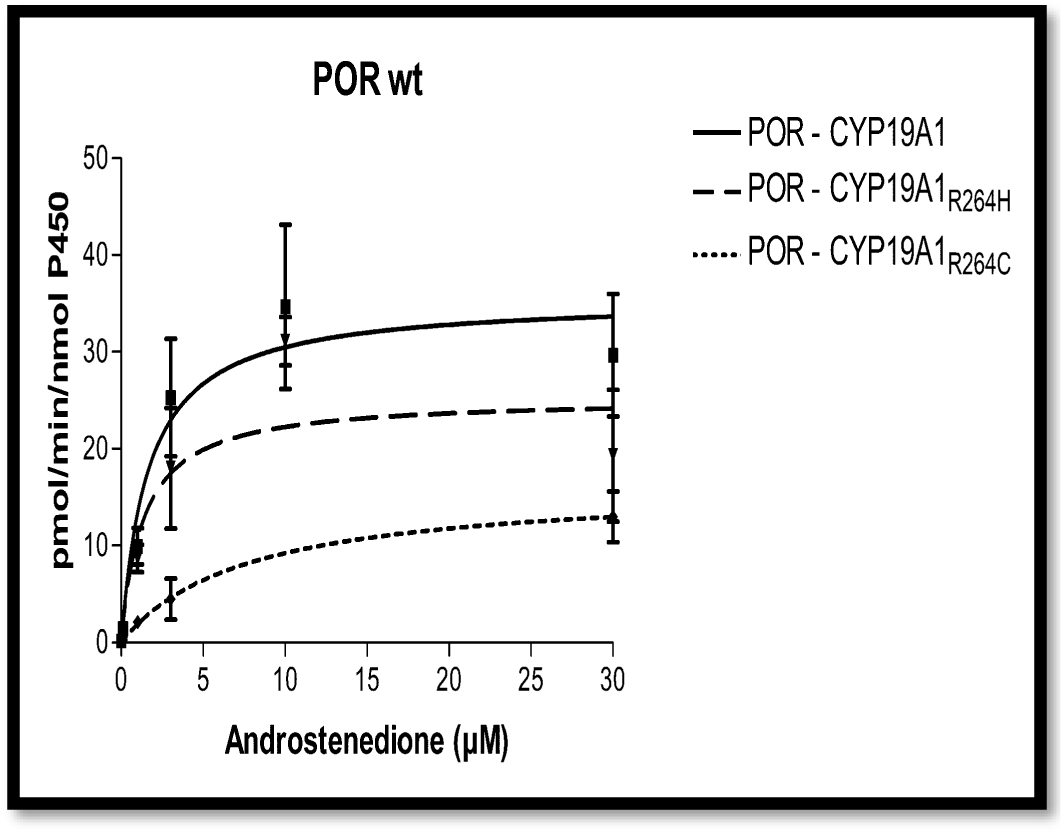
Aromatase activity of CYP19A1 variants with WT-POR. CYP19A1 wild type (solid line) and mutants CYP19A1_R264H_ (dashed line) and CYP19A1_R264C_ (dashed-dotted line) activities supported by POR-WT were tested to compare the activities of three different types of CYP19A1 variants. Recombinant preparations of purified human CYP19A1 (WT and variants) expressed in bacteria and recombinant human POR were combined into liposomes. CYP19A1 activity to biotransform androstenedione (^3^H labeled) to estrone was measured by monitoring the release of tritiated water. Data are shown as mean ± SEM of three independent replicates. Data were fitted using GraphPad Prism, according to the Michaelis-Menten kinetics model. The K_m_ and V_max_ values calculated from these assays and statistical analysis are given in Table 3.

Therefore, the CYP19A1-R264C variant showed a seven-fold reduction in catalytic efficiency (Vmax/Km) with only 15% of the residual CYP19A1 activity as compared to WT CYP19A1 (Figure 3, Table 3). This loss of catalytic efficiency indicates that R264C mutation of CYP19A1 affects either the interaction of the substrate with CYP19A1 or the CYP19A1-POR interaction, which may influence electron transfer from POR to CYP19A1 for metabolic reactions.

### Stability analysis of CYP19A1 and its polymorphic variants R264H and R264C by Fast Proteolysis Assay (FASTpp)

To compare the biophysical protein stability of CYP19A1 and its polymorphic variants R264H and R264C, we used a FASTpp assay with the recombinantly expressed and purified CYP19A1 proteins. CYP19A1-WT and variants were exposed in parallel to a range of different temperatures with the thermostable protease Thermolysin. Thermolysin readily cleaves the unfolded proteins near the hydrophobic residues Phe, Leu, Ile, and Val (42,43) but does not cleave the folded parts of the protein. Increasing the reaction temperature induces the thermal unfolding of proteins and the temperature-dependent changes in the degradation pattern represent the readout for the thermal stability of proteins. The SDS-PAGE analysis of the parallel FASTpp of CYP19A1-WT and variants (Figure 4A and B) showed that the R264H and R264C mutations do not display any major changes in protein stability. Therefore, it seems likely that the conformational changes induced by R264C and R264H variations do not significantly alter the structural stability of CYP19A1. However, the structural changes caused by R264C and R264H variations may alter the delicate balance of protein-protein interactions that are needed for the transfer of electrons from POR to CYP19A1.

**Figure 4:**
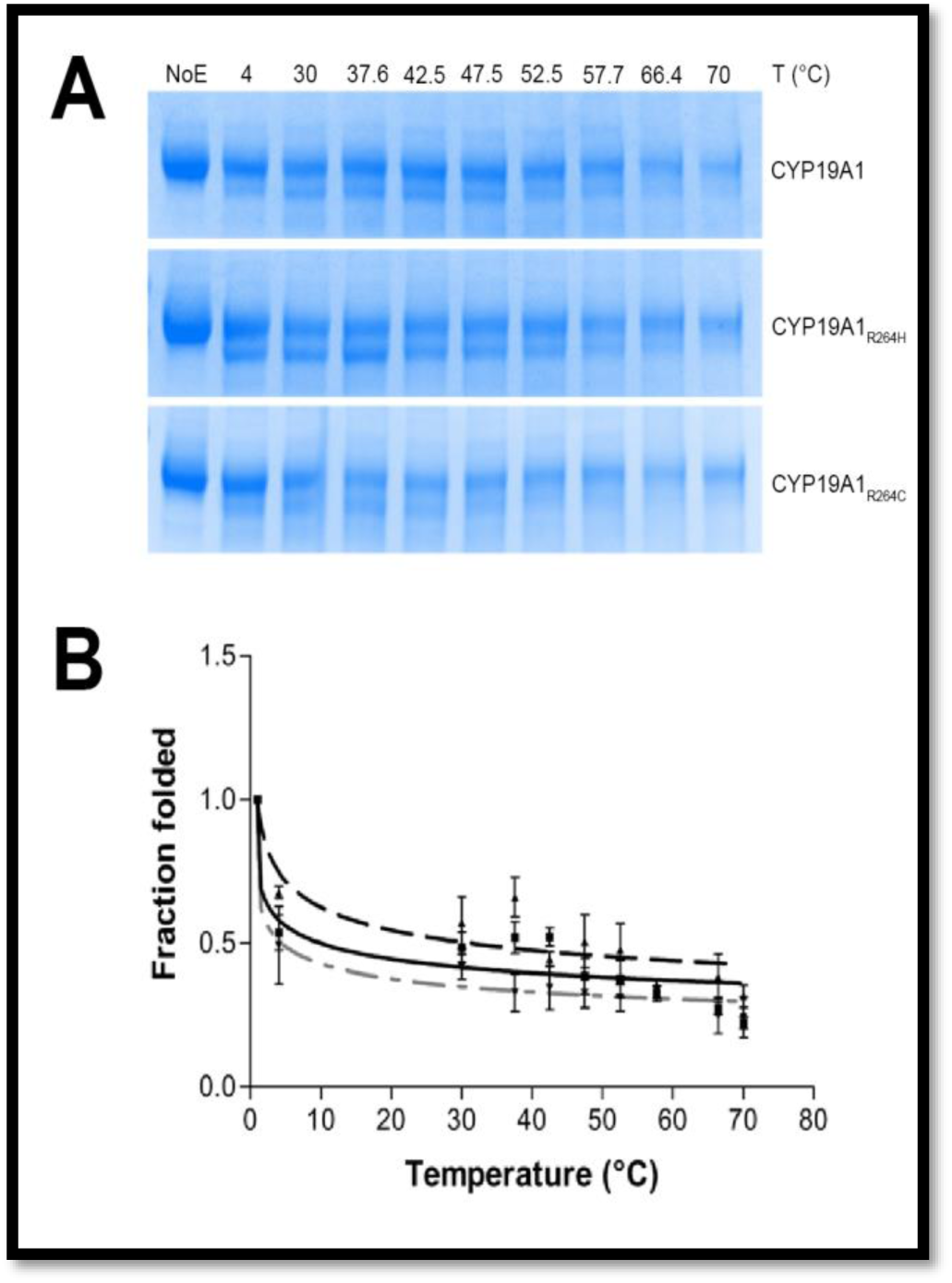
FastPP assay for the stability of CYP19A1 variants. Protein stability analysis of aromatase mutants CYP19A1_R264H_ (dashed line) and CYP19A1_R264C_ (dashed-dotted line) compared to CYP19A1-WT (solid line) by FASTpp assay. Thermolysin (0.05) mg/mL) was used to digest 0.20 mg/mL of CYP19A1-WT or mutants at increasing temperatures ranging from 4°C to 70°C. NoE represents the negative control carried out without thermolysin. The experiments were repeated three times, and here, a representative SDS-PAGE is shown. Each data-set represents mean ± SEM of three experiments. Data were analysed by Dunnett’s multiple comparisons test and the differences were not found to be statistically significant between the groups, indicating all variants have similar stability as the WT-CYP19A1.

### Variable effects on the enzyme activity of CYP19A1 and its polymorphic variants R264H and R264C are shown by POR variants

To study how different variants of CYP19A1 interact with variants of POR, we tested three polymorphic variants of POR (P284L, P284T, and Y607C) in supporting CYP19A1 enzyme activity. We tested the *in-vitro* estrogen synthase activity of CYP19A1 by the tritiated water release assay with the reconstituted liposomes using androstenedione as a substrate. The POR variants P284L and P284T showed a severe loss of enzyme activity with all CYP19A1 variants tested (Figure 5A-B, Table 4). For the POR variant P284L, the Km (apparent) for androstenedione was elevated ∼60 fold and ∼35 fold as compared to WT POR with CYP19A1 and CYP19A1-R264H. The apparent Vmax of POR-P284L was comparable to that of POR-WT in supporting CYP19A1 and CYP19A1-R264H activity. However, the product of enzymatic reaction with POR-P284L and CYP19A1-R264C was undetectable, and therefore, Km and Vmax could not be determined. For the POR-P284T, the Km (apparent) for androstenedione was elevated by ∼14, ∼32, and ∼12 fold with the CYP19A1-WT, CYP19A1-R264H, and CYP19A1-R264C respectively. The apparent Vmax was reduced by ∼55%, ∼30%, and ∼20% with the CYP19A1-WT, CYP19A1-R264H, and CYP19A1-R264C, respectively.

**Table 4.** Kinetic values for the activities of CYP19A1 and its polymorphic variants supported by variant forms of POR. For the conversion of androstenedione to estrone, variable amounts (0.3–30 µM) of androstenedione were used for enzyme assays, and the NADPH was fixed at 1mM. Data are shown as mean ± SEM of three independent replicates. Catalytic efficiencies of different combinations of POR and CYP19A1 variants are given as a percentage of WT-POR and WT-CYP19A1 combination (% of WT-WT catalytic efficiency). * represents statistically significant differences from the POR-WT control (*p<0.05 based on two tailed students t-test).

| | Km, Androstenedione<br>(μM) | Vmax<br>pmol/min/nmol | Vmax/Km | % of WT-WT<br>catalytic<br>efficiency |
| --- | --- | --- | --- | --- |
| ***POR-P284L*** |  |  |  |  |
| CYP19A1-WT | 99.7 ± 312.0 | 37.4 ± 92.7 | 0.4 | 1.7 |
| CYP19A1-R264H | 55.2 ± 49.7 | 38.6 ± 24.0 | 0.7 | 3.2 |
| CYP19A1-R264C | n.d. | n.d. | n.d. | n.d. |
| ***POR-P284T*** |  |  |  |  |
| CYP19A1-WT | 22.5 ± 36.2 | 15.8 ± 13.2 | 0.7 | 3.3 |
| CYP19A1-R264H | 51.0 ± 56.5 | 24.5 ± 18.7 | 0.5 | 2.2 |
| CYP19A1-R264C | 19.9 ± 31.7 | 28.7 ± 22.0 | 1.4 | 6.7 |
| ***POR-Y607C*** |  |  |  |  |
| CYP19A1-WT | 4.7 ± 2.5 | 18.3 ± 3.0* | 3.9* | 18.2* |
| CYP19A1-R264H | 1.3 ± 0.6 | 17.7 ± 1.7* | 13.6 | 63.1 |
| CYP19A1-R264C | 77.5 ± 133.6 | 81.1 ± 105.0 | 1.0 | 4.8 |

**Figure 5:**
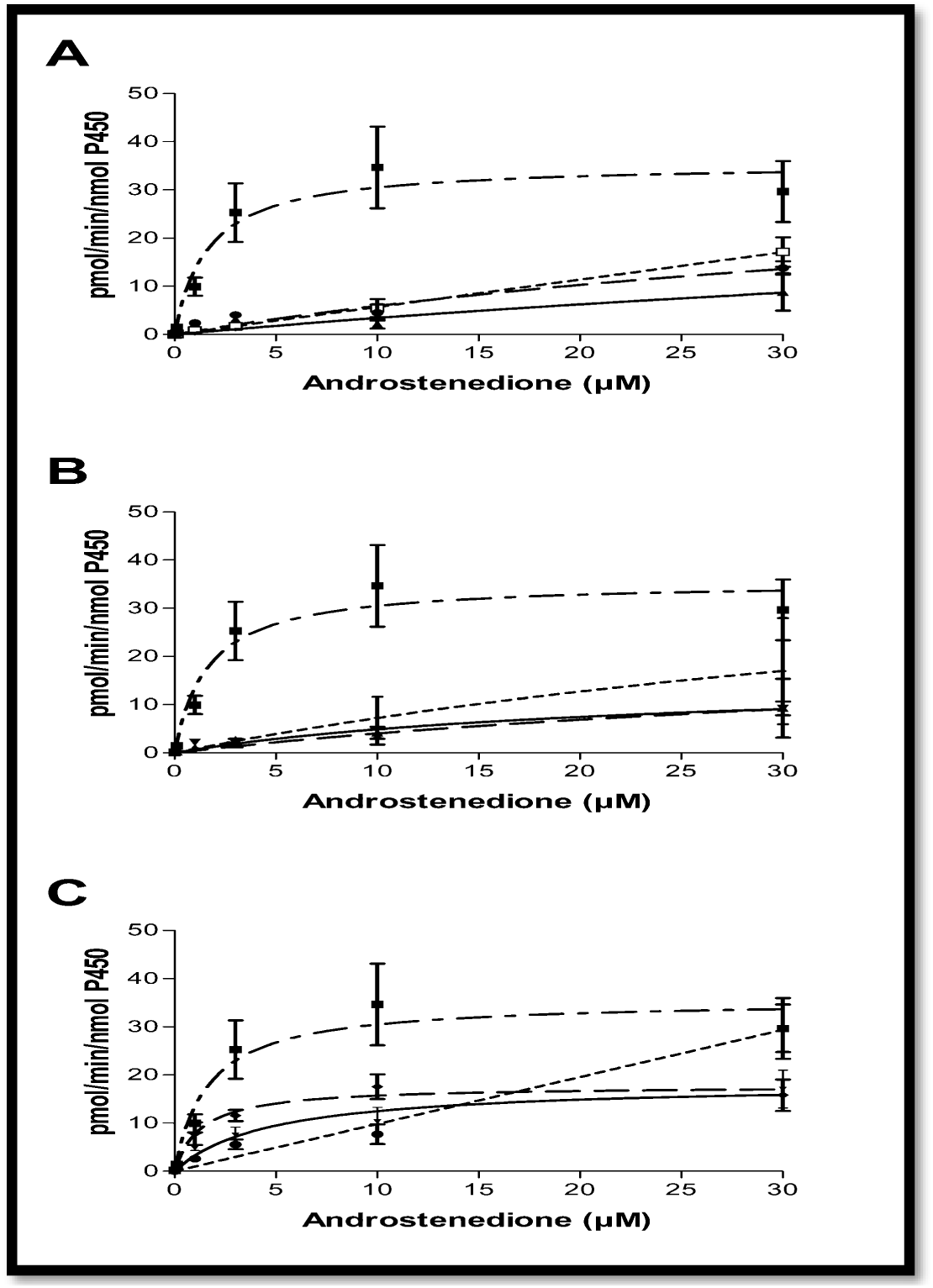
Enzyme assay of CYP19A1 variants. WT-CYP19A1(solid line), R264H-CYP19A1 (dashed line) and R264C-CYP19A1 (dotted line) activities supported by **A:** POR-P284L, **B:** POR-P284T, **C:** POR-Y607C. CYP19A1 activity supported by the wild type POR is shown in all three graphs as a reference (dash-dotted line). Human CYP19A1 (WT and variants R264C/R264H) and human POR (WT and variants P284L/P284T/Y607C) were embedded into liposomes, and aromatization activity to convert androstenedione into estrone was monitored by the release of ^3^H water during the reaction. Data are shown as mean ± SEM of three independent replicates. Data were fitted with GraphPad Prism using the kinetics model described by Michaelis and Menten. The K_m_ and V_max_ values calculated from these experiments and statistical analysis are shown in Table 3 and Table 4.

### POR-Y607C variant supports CYP19A1-R264H better than WT CYP19A1

For the POR variant Y607C, the Km (apparent) for androstenedione was elevated by ∼3 and ∼48 fold with the CYP19A1-WT and CYP19A1-R264C respectively (Figure 5C, Table 4). The apparent Vmax was reduced by ∼50%, with both CYP19A1-WT and CYP19A1-R264H. The residual catalytic efficiency of CYP19A1-WT, CYP19A1-R264H, and CYP19A1-R264C with POR-Y607C was 18.2%, 63.1 and 4.8% respectively as compared to POR-WT: CYP19A1-WT activity. Interestingly, the POR-Y607C variant supports the reaction of CYP19A1-R264H better than CYP19A1-WT.

## Discussion

POR is the obligate electron donor for all P450 enzymes in the endoplasmic reticulum and supplies electrons for the catalytic activities of CYP19A1 [42, 68, 84]. The CYP19A1 deficiency is linked to genetic disorders of estrogen biosynthesis in women and developmental disorders in men [19, 31, 46, 85, 86]. Previously we have tested the CYP19A1 variants (R264C and R26H) and POR genetic variants in separate studies that focused on individual genes [59, 64, 70]. Cytochromes P450 and P450 reductase have a large number of genetic variations and, therefore, many individuals may harbor variations in P450 reductase as well as P450 genes that may alter the enzymatic activities of cytochromes P450 [87]. So far, combined effects of variations in P450 reductase and steroidogenic P450 have not been tested, despite many reports on effects of genetic variations that were tested separately, using the WT P450 proteins with variant forms of POR; or P450 variants tested with WT-POR. There is only one report in literature describing the effects of genetic variations in a P450 protein (CYP2C9) with some POR variations [63].

In this report, we have tested the combinations of genetic variants in CYP19A1, responsible for estrogen production, with variant forms of POR. The R264C is a common polymorphism in CYP19A1 with a higher frequency in Asian and African populations compared to American, Mexican, or European populations. The only exception was found in the Maasai population from Kinyawa in Kenya, where the R264C variant of the CYP19A1 was present at a frequency similar to that of European or American populations (Table 1). The prevalence of the R264C variant of the CYP19A1 is high in Japanese, Chines, and southeast Asian populations. Among the POR variants studied, The Y607C has also been observed in the southeast Asian population [59]. In the CYP19A1 activity assays using bacterially expressed recombinant forms of POR and CYP19A1 proteins, we studied the combined effects of POR and CYP19A1 variants in different permutations and combinations of the two proteins. First, we tested the variant forms of CYP19A1 with WT POR and found that the CYP19A1-R264C variant lost around 85% of CYP19A1 catalytic efficiency, while the catalytic efficiency of CYP19A1-R264H variant was decreased by 14% when compared to WT-CYP19A1. These data are similar to the previous reports, where the two CYP19A1 polymorphic variants showed a decrease in the catalytic efficiency compared to the WT protein [51]. However, in this work, the amount of the activity loss was different, and this can be explained with a different experimental setup due to the incorporation of the enzymes in liposomes, the use of higher amounts of POR (4:1 *versus* 1:1 ratio) and the different assay of CYP19A1 activity used in the study.

POR is a membrane-bound protein that uses NADPH as a substrate for electron transfer. The binding of NADPH results in a conformational change that brings NAHPH close to the bound FAD in POR for the transfer of electrons. Afterward, additional changes in conformation cause a tightening of the POR structure, which brings the FAD closer to the FMN for the transfer of electrons from FAD to FMN. The P450 redox partners of POR interact with the FMN binding domain through ionic interactions involving lysine and arginine residues on the surface of P450s and the glutamate and aspartate residues located on the surface of POR [88-91]. Considering the importance of the hinge region of POR in both the intermolecular electron transfer as well as interactions with partner proteins, we studied the effects of variations in the hinge region of POR on the enzymatic activities of CYP19A1 variants. The main reason behind such sensitivity of CYP19A1 towards changes in POR may be due to its requirement of six pairs of electrons to complete the conversion of androgens (androstenedione, testosterone) to estrogens (estrone, estradiol). By this logic, it is possible that changes in the conformation of POR that affect interactions with CYP19A1, could, in turn, affect CYP19A1 activities. The P284L (rs 72557938) and P284T (rs72557937) variants are situated in the flexible hinge region of POR, which is important for movements of different domains, including the interactions between FMN and FAD-binding domains for the transfer of electrons from FAD to FMN (Figure 2B). In a study by Miller laboratory which analyzed genetic variations of POR in different populations, the POR variant P284L was found with an allele frequency of 0.003 in the Chinese Americans [76], and in the ExAc dataset from NCBI, the P284L variant of POR has an allele frequency of 0.00004. Huang et al. have reported a 54% loss of CYP17A1 17α-hydroxylase activity with POR P284L but found the 17,20 lyase activity of CYP17A1 to be similar to the wild type POR [76]. Recently we have shown that the P284L and P284T variants of POR showed reduced enzymatic activities with several drug-metabolizing cytochromes P450 [83, 87]. There was a 55% loss of activity in CYP2C19 and CYP3A5 assays, while a 78% loss of activity was observed in CYP2C9 assays with the P284L variant of POR compared to the WT protein [87]. Flavin content was not affected by the variation P284L in POR [64], indicating a change in protein-protein interactions, both the inter as well as intramolecular; may be behind the reduced CYP19A1 activities. In case of P284T variant of POR a reduction of both the FAD as well as FMN content has been observed [83]. The drug metabolizing cytochrome P450 enzyme activities were also reduced when the POR-P284T was used as a redox partner (CYP2C9, 26%; CYP2C19, 44%; CYP3A4, 23% and CYP3A5, 44% of WT POR) [83]. Because the flexible hinge region of POR is necessary for the movement of domains that brings the FAD closer to the FMN for the transfer of electrons, a change in the hinge region may alter the electron transfer process in POR, affecting the activities of redox partners [92]. It is likely that some electron transfer partners of POR may show a more significant effects on their enzymatic activities, while other redox partners of POR might remain unaffected depending on the precise nature of their interactions with the POR [93].

As seen with the P284L, P284T, and the Y607C mutations, there may be variations of POR in the seemingly non-clinical populations that show adverse effects on the enzymatic activities of multiple cytochrome P450 enzymes. Since variations in POR are now linked to the changes in the metabolism of many drugs, steroids, and small molecule substrates, variations in the POR gene should be considered as a candidate for genetic defects causing abnormal steroid or drug metabolism [59, 60, 62, 64, 75]. Further studies on genetic variants of POR shall be conducted to check the potentially harmful effect of individual POR variants.

## Acknowledgments

This work was supported by grants to AVP from the Novartis Foundation for Medical-Biological Research (18A053), the Swiss National Science Foundation (31003A-134926), and Burgergemiende Bern and by a CRT Foundation grant (project “Exposome” RF = 2016.2780) to GDN.

